# Silver-spoon upbringing improves early-life fitness but promotes reproductive ageing in a wild bird

**DOI:** 10.1101/535625

**Authors:** Foteini Spagopoulou, Celine Teplitsky, Martin I. Lind, Lars Gustafsson, Alexei A. Maklakov

## Abstract

Early-life conditions can have long-lasting effects and organisms that experience a poor start in life are often expected to age at a faster rate. Alternatively, individuals raised in high-quality environments can overinvest in early-reproduction resulting in rapid ageing. Here we use a long-term experimental manipulation of early-life conditions in a natural population of collared flycatchers (*Ficedula albicollis*), to show that females raised in a low-competition environment (artificially reduced broods) have higher early-life reproduction but lower late-life reproduction than females raised in high-competition environment (artificially increased broods). Reproductive success of high-competition females peaked in late-life, when low-competition females were already in steep reproductive decline and suffered from a higher mortality rate. Our results demonstrate that “silver-spoon” natal conditions increase female early-life performance at the cost of faster reproductive ageing and increased late-life mortality. These findings demonstrate experimentally that natal environment shapes individual variation in reproductive and demographic ageing in nature.

## Introduction

Ageing, a physiological deterioration with advancing age, is a complex and highly variable trait that is omnipresent across the tree of life^1,2^ and affects organismal fitness in nature^3^. Theory maintains that ageing evolves because selection gradients on traits decline with age^4–6^, favouring alleles that increase early-life fitness even at the cost of negative pleiotropic fitness effects in late-life. While a large body of evidence supports these theoretical predictions on the population level, very little is known about the individual variation in ageing rates. There is a general agreement in the field that early-life environmental conditions can shape the life-history of an adult organism and explain much of the individual variation in ageing rates^7–9^, but there is little consensus regarding the direction of the effect. Good environmental conditions during development can result in lifelong positive effects on physiology, reproduction and longevity^9–13^ (“silver-spoon” hypothesis^14^). Alternatively, organisms raised in good natal environments can invest heavily in growth and early-life reproduction, resulting in accelerated senescence in late-life^8,15–17^ (“live-fast die-young” hypothesis). These studies suggest that organisms can strategically allocate resources to reproduction and somatic maintenance depending on their early-life environmental conditions, and that high latent costs of increased early-life growth and reproduction, under favourable conditions, can result in accelerated ageing^17,18^.

Previous studies emphasised the role of sexual selection in the evolution of resource allocation strategies that can result in rapid ageing under favourable natal conditions. Males, for example, are predicted to invest in costly secondary sexual traits and risky behaviours, trading-off with investment in somatic maintenance and late-life performance^12,15–17^. However, recent studies also show that early-life investment in reproduction trades-off with late-life fitness in females in natural populations [reviewed in ^19^] and in the laboratory^20^. This suggests that “silver-spoon” females could allocate more resources to early-life reproduction and suffer from faster senecence later in life, while females experiencing early-life stress could reduce their investment in early-life reproduction and delay senescence.

Interestingly, mild stress in early-life results in reduced early but improved late breeding performance in captive female Zebra Finches (*Taeniopygia guttata*)^21^. However, whether females do indeed make similar reproductive decisions under natural conditions remains largely unknown.

We studied the long-lasting effects of natal environment on age-specific survival and reproduction in female Collared Flycatchers (*Ficedula albicollis*) from a natural population on the Swedish island of Gotland. We experimentally manipulated the brood size of hundreds of nesting pairs across 26 breeding seasons (i.e. 1983-2009) to create either artificially increased (high-competition) or artificially reduced (low-competition) broods by two offspring compared to the original brood, as well as control broods, where two offspring were exchanged between nests. Individuals from nests with a reduced brood size experience a “silver-spoon” upbringing and grow to a larger body size at fledgling^22^, allowing us to test directly whether experimentally improved developmental conditions decelerate or accelerate reproductive and demographic ageing. To our knowledge, this is the first study that directly investigates the consequences of individual heterogeneity in early-life condition for ageing in an experimentally manipulated natural population.

**Figure 1.**
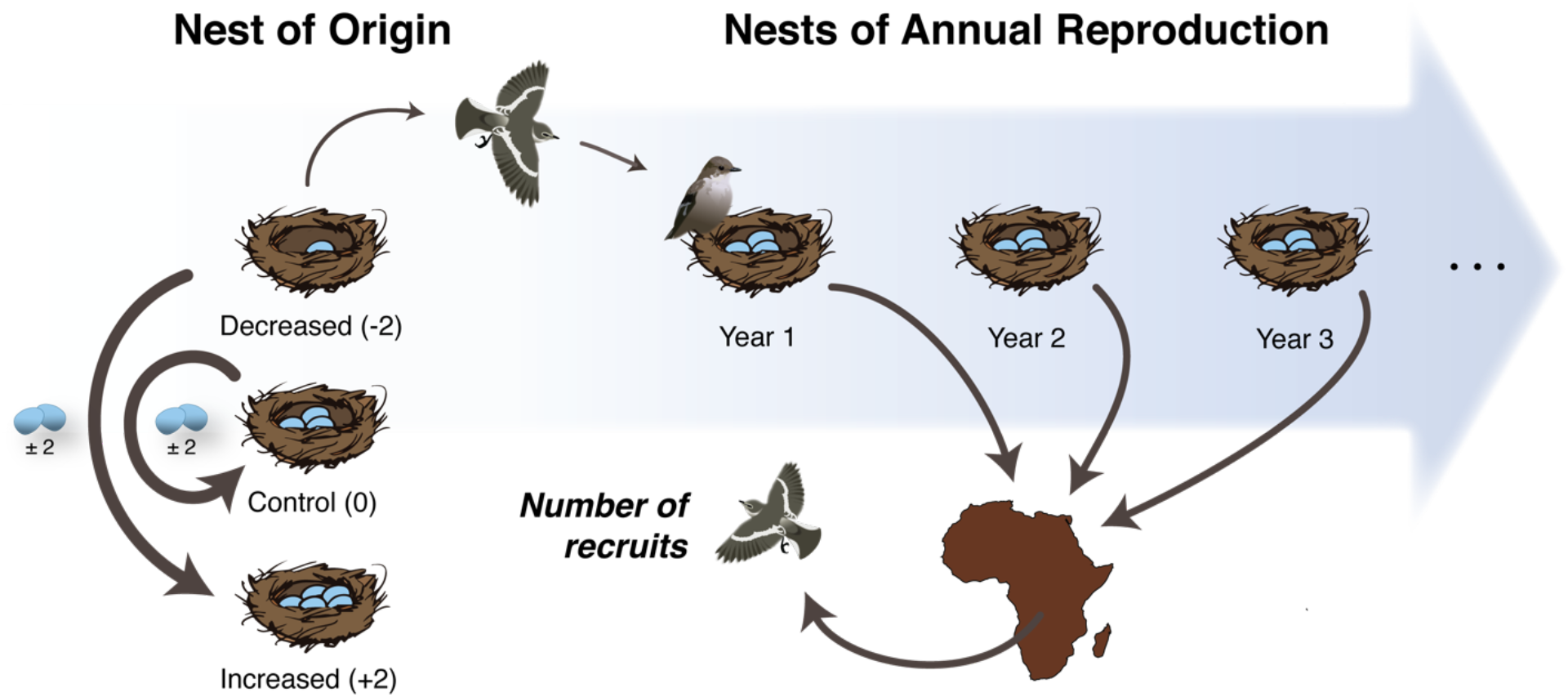
Experimental design for each of the 512 female collared flycatchers. The experimental brood size manipulation of the nest of origin is illustrated on the left and the age-specific reproductive data (i.e. number of recruits) collected from the nests of annual reproduction are illustrated on the right.

## Results

Experimental reduction of the brood size resulted in increased body mass at fledgling (15.12 ± 0.12, *χ*^2^ = 10.69, *P* < 0.01; Fig. 2A), while there was no effect of body size measured as tarsus length (19.56 ± 0.05, *χ*^2^ = 0.52, *P* = 0.77; Fig. S1). This suggests that brood reduction indeed improved natal environment and body condition of the offspring, rather than their structural size. Earlier studies showed that experimental increase of brood size results in reduced body mass at fledgling and increases offspring mortality^23–26^. Indeed, the body mass at fledgling of all the birds raised in our increased nests of origin is reduced (14.33 ± 0.11, *χ*^2^ = 36.35, *P* < 0.01; Fig. 2B). However, such an effect on the body mass of the focal high-competition treatment females is absent, due to the differential mortality within nests, which resulted in a subset of individuals surviving to reproduction.

**Figure 2.**
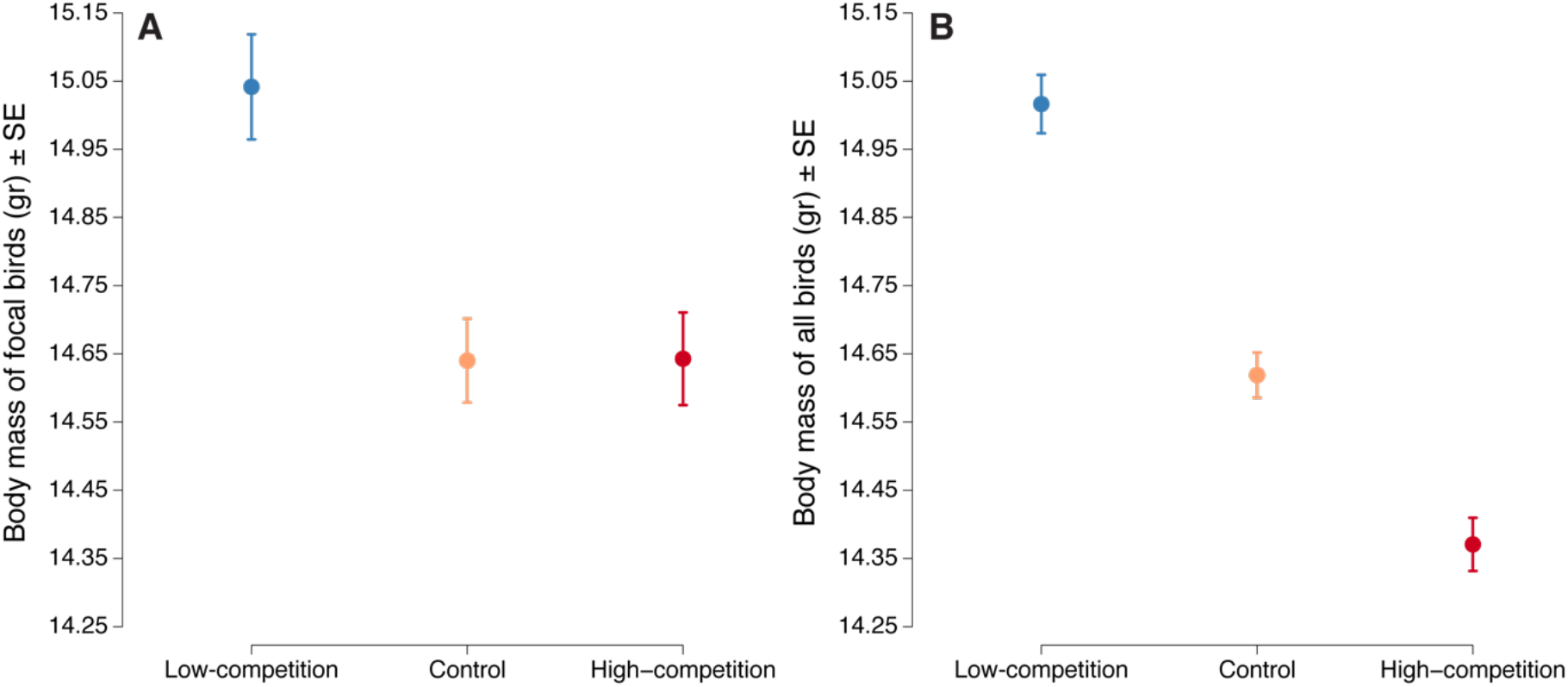
A) Body mass (means ± SE) of the focal birds raised in the three manipulated natal environments. B) Body mass (means ± SE) of all the birds raised in the three manipulated natal environments. Blue symbols represent the increased low-competition treatment, orange symbols represent the control treatment and red symbols represent the decreased high-competition treatment.

Even though, the quality of the natal environment had no significant effect on lifetime reproduction (*χ*^2^ = 0.90, *P* = 0.635; Fig. 3C, Table S1) or individual fitness *λ*_ind_ (*χ*^2^ = 2.17, *P* = 0.337; Fig. 3D, Table S2) in this study, females from the low-competition treatment had high reproductive success during the early years of their life, reaching a peak in the second and third year, and decline in the fourth (Fig. 3A-B, Table 1). In contrast, females raised in the high-competition treatment started low, but steadily increased their reproductive performance and peaked at years three and four (Fig. 3A-B, Table 1). While the raw data suggest that individuals from high-competition nests maintained high reproductive performance in their fifth year (Fig. 3A), taking into account selective appearance (measured as age at first reproduction) and disappearance (measured as age at last reproduction) reveals a reproductive decline in year five (Fig. 3B). Nevertheless, high-competition females showed a delayed reproductive peak and started to senescence later than their counterparts from low-competition nests. Females raised in control nests started at intermediate level of reproduction, maintained steady reproductive performance during the first three years of their life and declined at the fourth year.

**Figure 3.**
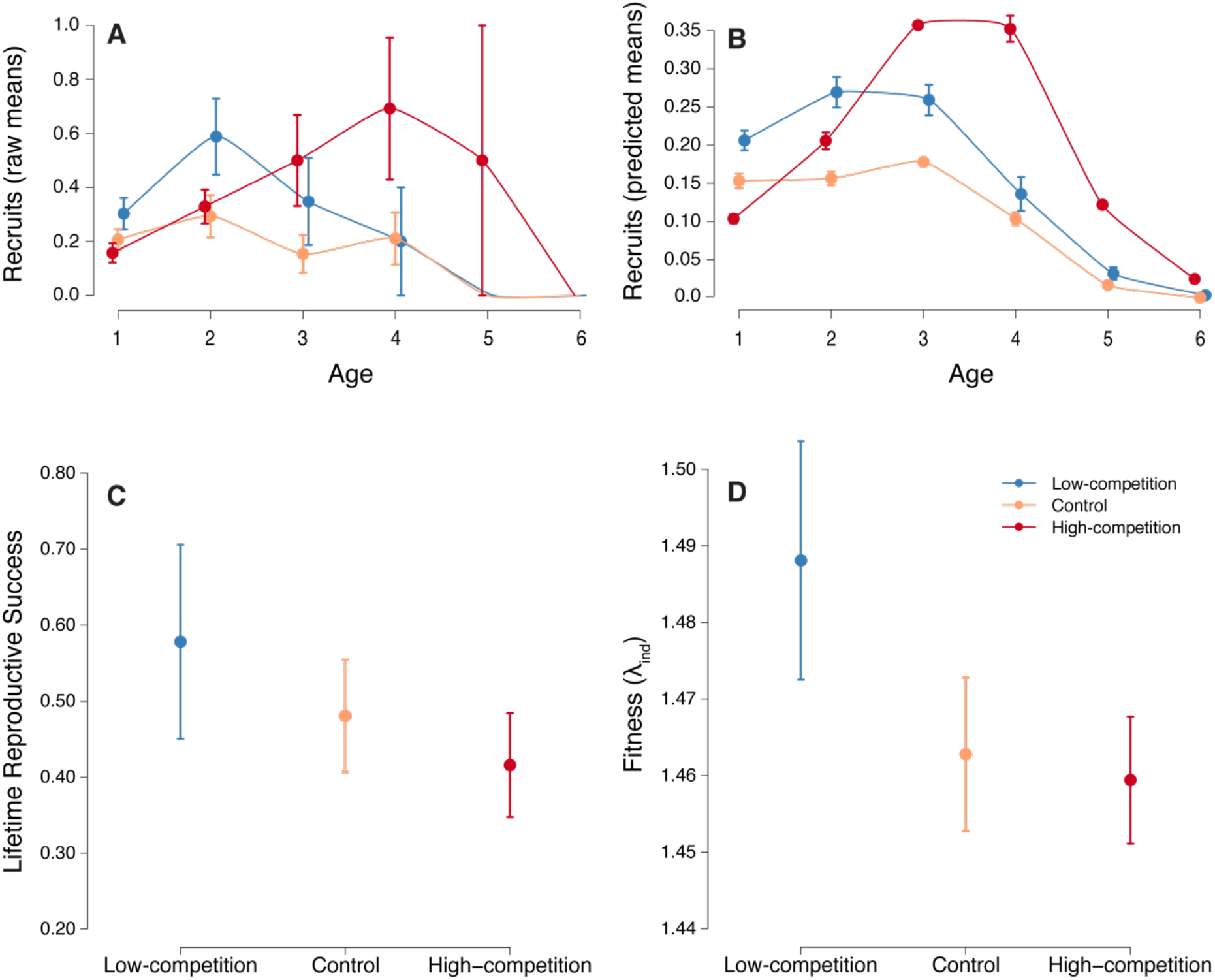
Reproduction of the three manipulated natal environments (Means ± SE). A) Age-specific reproduction from the raw data; B) Age-specific reproduction from the model predictions based on all random and fixed effects included in model 17 (Table 1); C) Lifetime Reproductive Success; D) Individual fitness (*λ*_ind_). Blue symbols represent the increased low-competition treatment, orange symbols represent the control treatment and red symbols represent the decreased high-competition treatment.

**Table 1.** Generalized linear mixed effects model examining the effect of early life environmental conditions on age-specific reproduction in 512 females. The reference level for treatment is “Low-competition”. Significant effects are in bold.

| Variable | Estimate<br>± S.E. | Z | P-value<br>(Z) | $\chi^2$ | df | P-value<br>($\chi^2$) | Variance |
| --- | --- | --- | --- | --- | --- | --- | --- |
| *Model 17* |  |  |  |  |  |  |  |
| **Intercept** | **-1.63 ± 0.26** | **-6.30** | **< 0.001** | **39.71** | **1** | **< 0.001** |  |
| **Age** | **0.62 ± 0.27** | **2.27** | **0.023** | **5.17** | **1** | **0.023** |  |
| **Age<sup>2</sup>** | **-0.52 ± 0.14** | **-3.64** | **< 0.001** | **13.24** | **1** | **< 0.001** |  |
| **Age at First Repr. (AFR)** | **-0.84 ± 0.16** | **-5.25** | **< 0.001** | **27.60** | **1** | **< 0.001** |  |
| Age at Last Repr. (ALR) | 0.31 ± 0.19 | 1.62 | 0.106 | 2.62 | 1 | 0.106 |  |
| Nest Treatment |  |  |  | 2.97 | 2 | 0.227 |  |
| Control Trt | -0.31 ± 0.22 | -1.43 | 0.150 |  |  |  |  |
| High-competition Trt | -0.33 ± 0.21 | -1.55 | 0.120 |  |  |  |  |
| Interaction (Age × ALR) | 0.14 ± 0.13 | 1.09 | 0.275 |  |  |  |  |
| **Interaction (Age × AFR)** | **0.92 ± 0.24** | **3.86** | **< 0.001** | **14.92** | **1** | **< 0.001** |  |
| Interaction (Age<sup>2</sup> × AFR) | -0.21 ± 0.11 | -1.84 | 0.065 | 3.39 | 1 | 0.065 |  |
| **Interaction (Age × Trt)** |  |  |  | **7.87** | **2** | **0.019** |  |
| Age × Control Trt | 0.24 ± 0.30 | 0.79 | 0.426 |  |  |  |  |
| **Age × High-competition Trt** | **0.79 ± 0.29** | **2.67** | **0.007** |  |  |  |  |
| Interaction (ALR × Trt) |  |  |  | 5.09 | 2 | 0.078 |  |
| **ALR × Control Trt** | **-0.55 ± 0.26** | **-2.08** | **0.037** |  |  |  |  |
| ALR × High-competition Trt | -0.47 ± 0.26 | -1.83 | 0.067 |  |  |  |  |
| Random effects |  |  |  |  |  |  |  |
| Female ID |  |  |  |  |  |  | 0.00 |
| Nest ID |  |  |  |  |  |  | 0.26 |
| Year of Birth |  |  |  |  |  |  | 0.00 |
| Year of Annual Repr. |  |  |  |  |  |  | 0.41 |
| Observation |  |  |  |  |  |  | 0.19 |

Natal environment also had profound effects on age-specific mortality rates (Fig. 4). Low-competition females had substantially lower initial acceleration in mortality rate (smaller early-life acceleration parameter b1 with KLDC > 0.85 – Fig. 4, Fig. 5 and Table S6) than both control and high-competition females, suggesting that “silver-spoon” natal environment is beneficial for survival. However, while the mortality rates of control and high-competition females strongly decelerated in late-life and reached a plateau around four years of age, the mortality rate of low-competition females continued to increase and reached a higher plateau around the fifth year (smaller late-life deceleration parameter *b*_2_ with KLDC > 0.85 – Fig. 4, Fig. 5 and Table S6).

**Figure 4.**
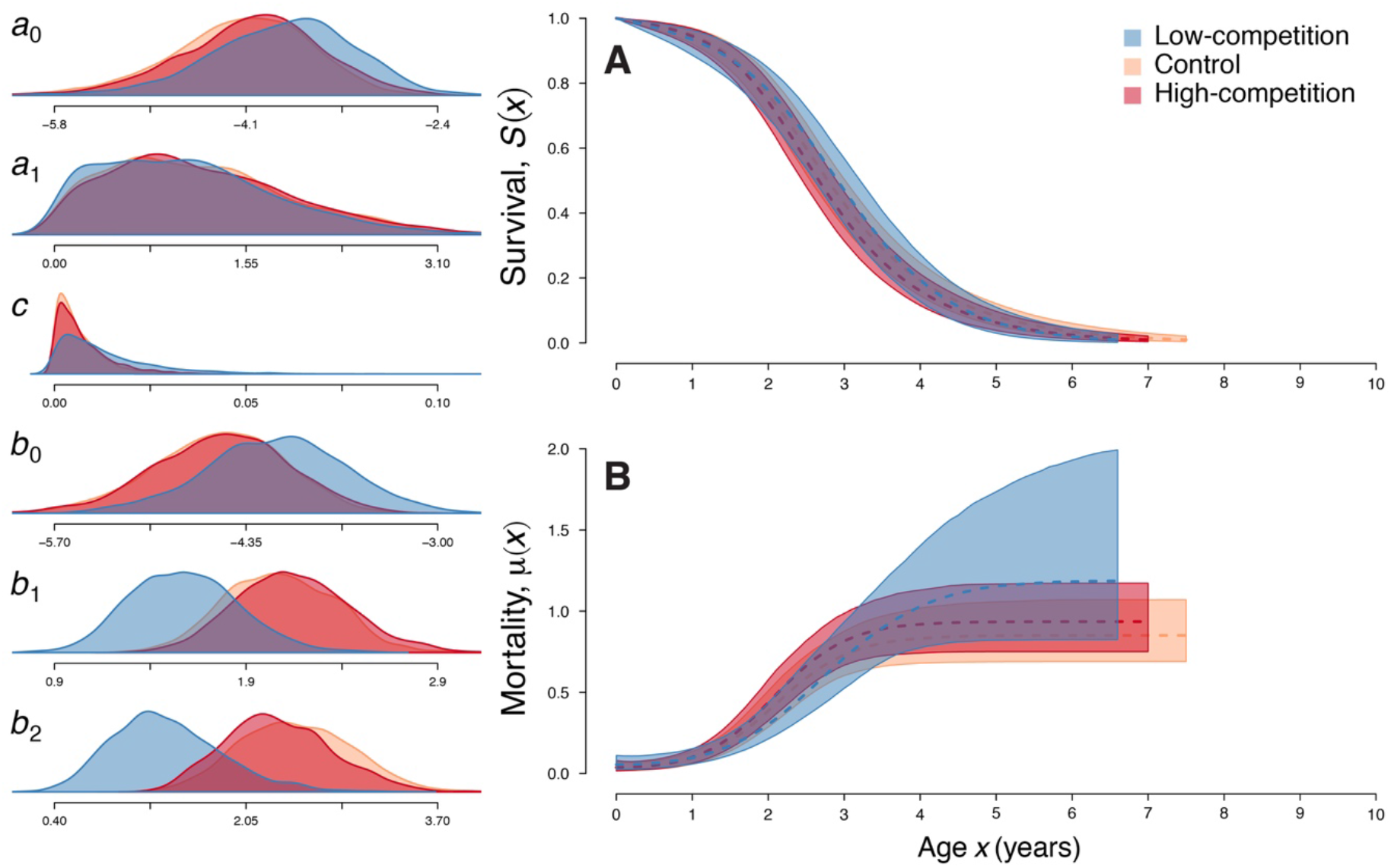
Survival (A) and Mortality (B) curves with 95% confidence intervals for each treatment as fitted by the bathtub Logistic mortality model. On the left are the posterior distributions of the six model parameters (see Methods for details). Blue colour and distributions represent the increased low-competition treatment, orange represent the control treatment and red represent the decreased high-competition treatment.

**Figure 5.**
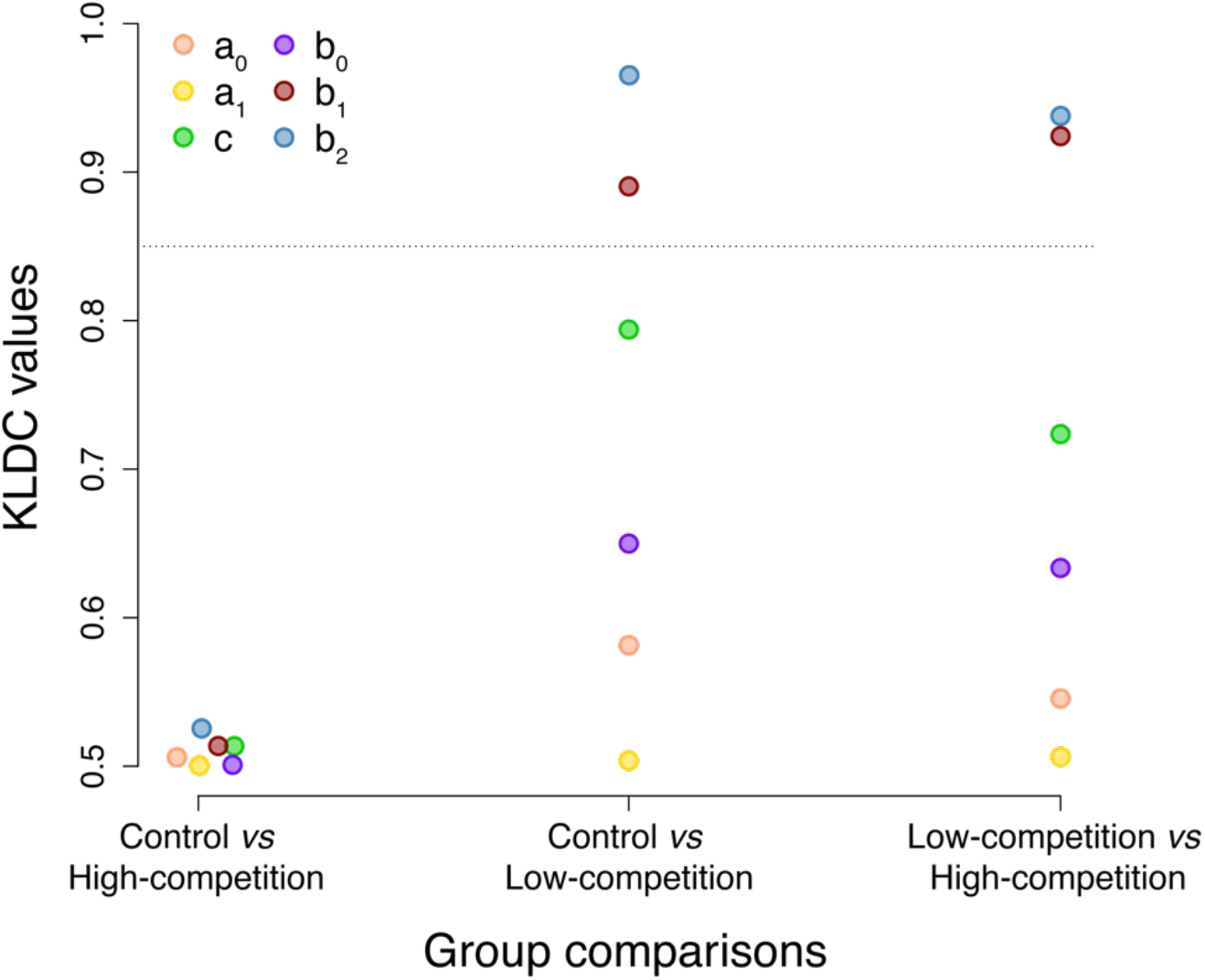
Values of the Kullback-Leibler divergence calibration (KLDC), which compare parameter posterior distributions of the bathtub Logistic mortality model, between the high-competition, the control and the low-competition treatments. The values that are above the dotted line (KLDC = 0.85) show parameters that are substantially different between the two treatments in each group comparison.

Thus, we found that females experiencing a low-competition natal environment fledge at higher body mass, start to reproduce at higher rate, have earlier reproductive peak and lower acceleration of mortality rate than their counterparts raised in high-competition nests. However, low-competition females also show earlier signs of reproductive senescence and suffer from increased mortality rate in the last years of their lives. This is the first experimental demonstration of condition-dependent ageing in a natural population. These results are support for the hypothesis that females raised in “silver-spoon” natal environment strategically allocate most of their resources to early-life reproduction, but pay a cost in terms of accelerated ageing in late-life.

## Discussion

There has been a sustained interest in how resource availability affects age-specific life histories but, remarkably, both theoretical and empirical studies to date are focused exclusively on the role of sexual selection in driving rapid ageing in high-condition males (i.e. “life fast-die young” hypothesis)^15–18,27–31^. This is, in part, because of the striking absence of studies showing that high-condition females may also trade-off early-life reproduction for accelerated ageing. Here, we used a long-term experimental manipulation of early-life environment to show that “silver-spoon” (i.e. low-competition) females indeed enjoy elevated early-life reproductive success, but suffer from earlier onset of reproductive ageing and increased mortality late in life. Thus, our findings call for a reappraisal of the idea that “live fast-die young” strategy applies primarily to males when faced with increased sexual competition^29,31^. Furthermore, our results suggest, that we should look beyond a simple negative correlation between reproduction and longevity when investigating the long-term fitness consequences of early-life conditions. Specifically, in this study “silver-spoon” females did not live shorter, nor reproduced more, but rather had different age-specific mortality and reproduction rates. It is possible that stochastic natural environment masks the effect of our brood size manipulation on total reproduction, and that the addition of several more years of data collection could show that high-condition females actually have higher fitness. At this point, however, we can only state that a “silver-spoon” early-life environment results in increased early-life but reduced late-life performance.

Our results are in line with the “disposable soma” theory of ageing, which maintains that early-life investment in growth and reproduction results in reduced late-life fitness due to an increased amount of unrepaired cellular damage^32–34^. However, such patterns can also emerge directly from increased molecular nutrient-sensing signalling in “silver-spoon” females that received more food during development, resulting in increased cellular hypertrophy and accelerated cellular senescence late in life^35,36^. Future research should focus on understanding the physiological and molecular underpinnings of the phenomenon that improved early-life conditions lead to detrimental effects in late-life.

## Methods

### Study species and study area

Collared flycatchers (*Ficedula albicollis*) are small (~ 13g), migratory, insectivorous, cavity-nesting passerines, that breed in deciduous and mixed forests of eastern and central Europe (and southwestern Asia). Females incubate four to eight eggs during a 14-day period and rear a single brood each year. Juveniles are fed by both parents and fledge 14 to 18 days after hatching (for further details on population and methods see ^37^).

Isolated from their main breeding grounds, additional breeding populations have been established on the Swedish islands of Gotland and Öland in the Baltic Sea.

The collared flycatcher population on the island of Gotland (57° 10°, 18° 20°) has been well studied since 1980. Each year, new individuals (i.e. both adults and offspring) are measured and ringed, to allow individual identification throughout their lifetime and enable the assessment of yearly reproductive success. The females are caught during incubation or the chick provisioning period and the males only during the chick provisioning period. Further details of the breeding region, and ecology have been collected using standard methods^37,38^.

Collared flycatchers in general and the long-term studied Gotland population in specific, are particularly suitable for the study of life history trade-offs in nature due to their high degree of site fidelity, limited dispersal and preference for nest boxes over natural cavities, which allows accurate age and survival estimation ^38^.

### Manipulations & data selection

Since 1983, multiple brood size manipulation experiments have been conducted in the Gotland population^26,39–41^, to examine various aspects of parental investment, the costs of increased reproductive effort and the trade-off between reproductive investment and immune function. To investigate the effects of early-life natal conditions on individual life histories, we collected data from individuals raised in manipulated nests, where clutch size was increased or reduced by two offspring. In addition, we accounted for possible translocation effects by including a control treatment, in which offspring were swapped between nests without alteration of the initial total number (Fig. 1). We monitored the entire life-history of 943 collared flycatchers, which were raised in such manipulated nests, but eventually focused on females, due to the degree of male extra-pair paternity (~15% ^42^) that makes the process of estimating male reproductive values challenging.

Reproductive and survival data were collected for 512 females born between 1983 and 2009. 121 females were raised in nests with reduced broods (“low competition”), 214 females were raised in controlled nests (“control”) and 177 females were raised in nests with increased broods (“high competition”). For each female, we recorded the number of recruits (i.e. offspring that returned to Gotland to breed the following years, after overwintering in Africa; Fig. 1) during each reproductive season and used this measurement to analyse how reproduction changes with increasing age (age-specific reproduction). The annual number of recruited offspring was then summed across all years and yielded an estimate of lifetime reproductive success; a major individual fitness component. Finally, we also analysed the reproductive data using the rate-sensitive individual fitness *λ*_ind_. This measurement encompasses the number of offspring, as well as their timing^43,44^. and is, thus, analogus to the intrinsic rate of population growth^45^.

Over the years, some of our focal females (N = 204) were involved in additional experiments addressing different questions^40,41,46^. To account for this in our analysis, we excluded the current and/or future reproductive values of each affected bird, depending on the nature of these additional experimental manipulations and the effect they may have on the female’s reproduction.

The age-specific reproductive patterns were similar in both the full (N = 512) and the restricted dataset (N = 308), where only the subset of individuals that were not subjected in such manipulations was included. For the lifetime reproductive analysis and the rate-sensitive individual fitness (*λ*_ind_) analysis, we only used the restricted dataset.

### Statistical analysis

All analyses were performed in *R* version 3.3.1^47^. Statistical models were fitted using the *lme4* package (v.1.1-14^48^) and the *MCMCglmm* package (v.2.23^49^). For generalized linear mixed models (Poisson error structure), we ensured an appropriate fit using the *DHARMa* package (v. 0.1.2) for diagnostic tests of model residuals^50^. We used the package *AICcmodavg* (v. 2.1-0^51^) to compute the Akaike’s Information Criterion for small sample sizes (AICc). To test for significance of model terms we used the Anova function of the *car* package (v. 2.1-3) with Type III Wald chi-square tests. Finally, for the survival analysis models we used the Bayesian framework from the *BaSTA* package (v. 1.9.5).

#### Reproductive success analysis

To analyse the effects of the brood size manipulation on the age-specific reproduction, we used both a frequentist and a Bayesian framework, which yielded the same results. Thus, we describe below the frequentist approach, but see Supplementary Methods for further details on the Bayesian approach.

We constructed a generalised linear mixed-effects model (GLMM) with Poisson error structure and log link transformation. We accounted for repeated measurements and cohort effects by including individual identity, nest of origin identity, year of origin and year of annual reproduction as random effects. Furthermore, we included an observation-level random effect to control for overdispersion^52^. As fixed effects, we included the treatment (increased, reduced and control broods), as well as a linear and a quadratic term of age (Age and Age^2^), to test for non-linear variation of reproductive value with age. In addition, we included the Age of First Reproduction (AFR) and Age at Last Reproduction (ALR) in our models, to control for selective appearance and selective disappearance^53^. All continuous predictor variables were standardised (mean = 0 and SD = 1), to allow an interpretation of relative effect sizes.

We initially run a complete model including all fixed effects and up to three-way interactions (Full model). We then used backward deletion of non-significant variables (P > 0.05) with all possible combinations in the order of terms we deleted, starting with the highest order interaction terms; a process that resulted in 35 different models. We then compared all models using the conservative Akaike Information Criterion for small sample sizes (AIC_c_). Three of these models had the smallest AIC_c_ values, indicating best model fit (models 17, 24 and 31; see Table S3). They all gave similar outcomes and we therefore only present here the most comprehensive model (model 17) that includes most variables, but see supplementary material for the output of the other two models (Suppl. Table S4).

For the lifetime reproduction, we constructed a GLMM with Poisson error structure and log link transformation, using the lifetime number of recruits as the response variable and treatment as fixed effect. We used nest of origin identity and year of birth as random effects to control for cohort effects (Table S1).

Finally, for the analysis of rate-sensitive individual fitness, we first estimated *λ*_ind_ by solving the Euler-Lotka equation of age-specific recruit number and survival for each individual using the *lamda* function in the *popbio* package (v 2.4.4^54^). Then we constructed a LMM model, using the log of *λ*_ind_ as the response variable. Similarly with the lifetime reproductive analysis, we included the treatment as a fixed effect. Finaly the nest of origin and year of birth were used as random factors to control for cohort effects (Table S2).

#### Survival analysis

To investigate how early-life developmental conditions affect survival and mortality rates of female collared flycatchers, we performed a Bayesian Survival Trajectory Analysis (BaSTA)^55^. BaSTA is a Bayesian approach that uses Markov Chain Monte Carlo (MCMC) procedures to optimize mortality distributions estimating the slope with which mortality increases with age (i.e. demographic ageing). It is a suitable package to fit mortality distributions and estimate parametric survival functions of capture-recapture data, like ours, where information about the exact age of death is missing^56^. It estimates the recapture rate and uses this probability of recapture to correct the survival estimates extracted from the BaSTA survival models. The recapture probability in our data was 69% (Table S6).

The survival analyses assume no dispersal, which could bias the effect of natal environment on actuarial senescence. Indeed, collared flycatchers show high site fidelity and very limited dispersal^38^. Furthermore, given that the year of birth is known for every individual in our analysis, we are therefore confident that the population estimates of the mortality distribution parameters are reliable^56^.

We explored actuarial mortality rates using Weibull, Gompertz and Logistic models, with either a simple shape or one of the more complex Makeham^57^ or bathtub^58^ shapes. Furthermore, we also explored a tenth model, the Exponential model with a simple shape. We performed four parallel simulations that each ran for 2 200 000 iterations, used a burn-in of 200 000 chains and took a model sample each 4 000 chains. The model parameters likely converged on the most optimal mode and showed low serial autocorrelations (<5%) and robust posterior distributions (with N= 2 000). For the model comparison, we used the lowest deviance information criterion (DIC), which is similar to the Akaike Information Criterion (AIC) in non-Bayesian models. Comparison of the DIC values^59^ revealed that the Logistic model with the bathtub shape is the most appropriate mortality function (Table S7). Mortality rate:

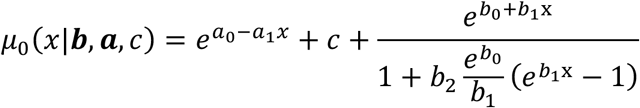

The bathtub structure used in the BaSTA package adds a declining Gompertz function and a constant to the basic logistic mortality function. Therefore, the two alpha parameters (α_0_ and α_1_) describe the exponential decline that may be observed very early in life and the constant (c) describe the lowest point of the mortality function. The beta parameters (b_0_, b_1_ and b_2_) describe different parts of the logistic increase in mortality rates with age (i.e. initial exponential increase in mortality that decelerates until it reaches a plateau). More specifically, b_0_ is the age-independent or baseline mortality, b_1_ describes the initial exponential increase in mortality with age and b_2_ describes the degree of deceleration in mortality with age and the level of the asymptote.

To assess the differences among the posterior distributions of our brood size manipulation treatments we used the Kullback-Leibler divergence calibration (KLDC)^60^, which show the mean Kullback-Leibler discrepancies (KL) between two distributions. Values closer to 0.5 imply that there is minimal difference among the distributions and values closer to 1 imply major differences^60,61^. We considered a KLDC value > 0.85 to indicate a substantial difference between the two posterior distributions that are compared (Fig. 5).

## Acknowledgments

We thank everyone involved in collecting the flycatcher data over the years. This work has been supported by the ERC Starting Grant AGINGSEXDIFF 260885 and the ERC Consolidator Grant GermlineAgeingSoma 724909 to A.A.M. The long-term field study have been supported continuously by grants from the Swedish Research Council (VR) to L.G.

## Statement of authorship

L.G. collected the data. A.A.M, L.G. and F.S conceived the project. F.S., C.T. and M.I.L. performed the reproductive data analysis. F.S performed the survival analysis. A.A.M. and L.G. contributed to data analysis and interpretation. F.S. and A.A.M wrote the manuscript. All authors contributed substantially to revisions.

## References

1. Jones, O. R. et al. Diversity of ageing across the tree of life. Nature 505, 169–173 (2014).

2. The evolution of senescence in the tree of life. (Cambridge University Press, 2017).

3. Bouwhuis, S., Choquet, R., Sheldon, B. C. & Verhulst, S. The forms and fitness cost of senescence: age-specific recapture, survival, reproduction, and reproductive value in a wild bird population. Am. Nat. 179, E15–E27 (2012).

4. Medawar, P. B. An unresolved problem of biology. (London: H.K. Lewis, 1952).

5. Williams, G. C. Pleiotropy, natural selection, and the evolution of senescence. Evolution 11, 398–411 (1957).

6. Hamilton, W. D. The moulding of senescence by natural selection. J. Theor. Biol. 12, 12–45 (1966).

7. Monaghan, P. Early growth conditions, phenotypic development and environmental change. Philos. Trans. R. Soc. Lond. B Biol. Sci. 363, 1635–1645 (2008).

8. Marshall, H. H. et al. Lifetime fitness consequences of early-life ecological hardship in a wild mammal population. Ecol. Evol. 7, 1712–1724 (2017).

9. Cooper, E. B. & Kruuk, L. E. B. Ageing with a silver-spoon: A meta-analysis of the effect of developmental environment on senescence. Evol. Lett. 2, 460–471 (2018).

10. Lindström, J. Early development and fitness in birds and mammals. Trends Ecol. Evol. 14, (1999).

11. van de Pol, M., Bruinzeel, L. W., Heg, D., Van Der Jeugd, H. P. & Verhulst, S. A silver spoon for a golden future: long-term effects of natal origin on fitness prospects of oystercatchers (*Haematopus ostralegus*). J. Anim. Ecol. 75, 616–626 (2006).

12. Nussey, D. H., Kruuk, L. E. B., Morris, A. & Clutton-Brock, T. H. Environmental conditions in early life influence ageing rates in a wild population of red deer. Curr. Biol. 17, R1000–R1001 (2007).

13. Hayward, A. D., Rickard, I. J. & Lummaa, V. Influence of early-life nutrition on mortality and reproductive success during a subsequent famine in a preindustrial population. Proc. Natl. Acad. Sci. 110, 13886–13891 (2013).

14. Grafen, A. Biological signals as handicaps. J. Theor. Biol. 144, 517–546 (1990).

15. Hunt, J. et al. High-quality male field crickets invest heavily in sexual display but die young. Nature 432, 1024–1027 (2004).

16. Bonduriansky, R. & Brassil, C. E. Reproductive ageing and sexual selection on male body size in a wild population of antler flies *(Protopiophila litigata):* Ageing and sexual selection. J. Evol. Biol. 18, 1332–1340 (2005).

17. Hooper, A. K., Spagopoulou, F., Wylde, Z., Maklakov, A. A. & Bonduriansky, R. Ontogenetic timing as a condition-dependent life history trait: High-condition males develop quickly, peak early, and age fast. Evolution 71, 671–685 (2017).

18. Adler, M. I., Telford, M. & Bonduriansky, R. Phenotypes optimized for early-life reproduction exhibit faster somatic deterioration with age, revealing a latent cost of high condition. J. Evol. Biol. 29, 2436–2446 (2016).

19. Lemaître, J.-F. et al. Early-late life trade-offs and the evolution of ageing in the wild. Proc. R. Soc. Lond. B Biol. Sci. 282, 20150209 (2015).

20. Travers, L. M., Garcia-Gonzalez, F. & Simmons, L. W. Live fast die young life history in females: evolutionary trade-off between early life mating and lifespan in female *Drosophila melanogaster*. Sci. Rep. 5, 15469 (2015).

21. Marasco, V., Boner, W., Griffiths, K., Heidinger, B. & Monaghan, P. Environmental conditions shape the temporal pattern of investment in reproduction and survival. Proc. R. Soc. B Biol. Sci. 285, 20172442 (2018).

22. Voillemot, M. et al. Effects of brood size manipulation and common origin on phenotype and telomere length in nestling collared flycatchers. BMC Ecol. 12, 17 (2012).

23. Alatalo, R. V., Gustafsson, L. & Lundberg, A. Phenotypic Selection on Heritable Size Traits: Environmental Variance and Genetic Response. Am. Nat. 135, 464–471 (1990).

24. Tinbergen, J. M. & Boerlijst, M. C. Nestling weight and survival in individual great tits (*Parus major*). J. Anim. Ecol. 59, 1113–1127 (1990).

25. de Kogel, C. H. Long-term effects of brood size manipulation on morphological development and sex-specific mortality of offspring. J. Anim. Ecol. 66, 167–178 (1997).

26. Sendecka, J., Cichoń, M. & Gustafsson, L. Age-dependent reproductive costs and the role of breeding skills in the Collared flycatcher. Acta Zool. 88, 95–100 (2007).

27. Kokko, H. Good genes, old age and life-history trade-offs. Evol. Ecol. 12, 739–750 (1998).

28. Radwan, J. & Bogacz, I. Comparison of life-history traits of the two male morphs of the bulb mite, Rhizoglyphus robini. Exp. Appl. Acarol. 24, 115–121 (2000).

29. Bonduriansky, R., Maklakov, A., Zajitschek, F. & Brooks, R. Sexual selection, sexual conflict and the evolution of ageing and life span. Funct. Ecol. 22, 443–453 (2008).

30. Preston, B. T., Jalme, M. S., Hingrat, Y., Lacroix, F. & Sorci, G. Sexually extravagant males age more rapidly. Ecol. Lett. 14, 1017–1024 (2011).

31. Hooper, A. K., Lehtonen, J., Schwanz, L. E. & Bonduriansky, R. Sexual competition and the evolution of condition-dependent ageing. Evol. Lett. 2, 37–48 (2018).

32. Kirkwood, T. B. L. & Holliday, R. The evolution of ageing and longevity. Proc. R. Soc. Lond. B Biol. Sci. 205, 531–546 (1979).

33. Kirkwood, T. B. L. Evolution of ageing. Mech. Ageing Dev. 123, 737–745 (2002).

34. Kirkwood, T. B. L. The Disposable Soma Theory. in The Evolution of Senescence in the Tree of Life (eds. Jones, O. R., Shefferson, R. P. & Salguero-Gómez, R.) 23–39 (Cambridge University Press, 2017). doi:10.1017/9781139939867.002

35. Blagosklonny, M. V. Aging and immortality: Quasi-programmed senescence and its pharmacologic inhibition. Cell Cycle 5, 2087–2102 (2006).

36. Gems, D. & Partridge, L. Genetics of longevity in model organisms: Debates and paradigm shifts. Annu. Rev. Physiol. 75, 621–644 (2013).

37. Gustafsson, L. Collared flycatcher. in Lifetime reproduction in birds (ed. Newton, I.) 75–88 (London: Academic Press, 1989).

38. Pärt, T. & Gustafsson, L. Breeding dispersal in the collared flycatcher *(Ficedula albicollis):* Possible causes and reproductive consequences. J. Anim. Ecol. 58, 305–320 (1989).

39. Gustafsson, L. & Sutherland, W. J. The costs of reproduction in the collared flycatcher *Ficedula albicollis*. Nature 335, 813–815 (1988).

40. Nordling, D., Andersson, M., Zohari, S. & Gustafsson, L. Reproductive effort reduces specific immune response and parasite resistance. Proc. R. Soc. Lond. B Biol. Sci. 265, 1291–1298 (1998).

41. Doligez, B. et al. Costs of reproduction: assessing responses to brood size manipulation on life-history and behavioural traits using multi-state capture-recapture models. J. Appl. Stat. 29, 407–423 (2002).

42. Sheldon, B. C. & Ellegren, H. Sexual selection resulting from extrapair paternity in collared flycatchers. Anim. Behav. 57, 285–298 (1999).

43. Brommer, J. E., Merilä, J. & Kokko, H. Reproductive timing and individual fitness. Ecol. Lett. 5, 802–810 (2002).

44. Lind, M. I., Zwoinska, M. K., Meurling, S., Carlsson, H. & Maklakov, A. A. Sex-specific Tradeoffs With Growth and Fitness Following Life-span Extension by Rapamycin in an Outcrossing Nematode, *Caenorhabditis remanei*. J. Gerontol. Ser. A 71, 882–890 (2016).

45. Stearns, S. C. The evolution of life histories. (Oxford University Press, 1992).

46. Pitala, N. et al. The effects of experimentally manipulated yolk androgens on growth and immune function of male and female nestling collared flycatchers *Ficedula albicollis*. J. Avian Biol. 40, 225–230 (2009).

47. R Development Core Team. R: A language and environment for statistical computing. R Found. Stat. Comput. Vienna Austria (2017).

48. Bates, D., Mächler, M., Bolker, B. & Walker, S. Fitting linear mixed-effects models using lme4. J. Stat. Softw. 67, 1–48 (2015).

49. Hadfield, J. D. MCMC Methods for multi-response generalized linear mixed models: The MCMCglmm R package. J. Stat. Softw. 33, 1–22 (2010).

50. Hartig, F. DHARMa: residual diagnostics for hierarchical (multi-level/mixed) regression models. R Package Version 010 (2016).

51. Mazerolle, M. J. AICcmodavg: Model selection and multimodel inference based on (Q)AIC(c). R Package Version 21–0 (2016).

52. Harrison, X. A. Using observation-level random effects to model overdispersion in count data in ecology and evolution. PeerJ 2, e616 (2014).

53. van de Pol, M., Verhulst, S., Pfister, A. E. C. A. & DeAngelis, E. D. L. Age-dependent traits: A new statistical model to separate within- and between-individual effects. Am. Nat. 167, 766–773 (2006).

54. Stubben, C. & Milligan, B. Estimating and Analyzing Demographic Models Using the popbio Package in R. J. Stat. Softw. 22, (2007).

55. Colchero, F., Jones, O. R. & Rebke, M. BaSTA: an R package for Bayesian estimation of age-specific survival from incomplete mark-recapture/recovery data with covariates: *BaSTA - Bayesian Survival Trajectory Analysis*. Methods Ecol. Evol. 3, 466–470 (2012).

56. Colchero, F. & Clark, J. S. Bayesian inference on age-specific survival for censored and truncated data. J. Anim. Ecol. 81, 139–149 (2012).

57. Pletcher, S. D. Model fitting and hypothesis testing for age-specific mortality data. J. Evol. Biol. 12, 430–439 (1999).

58. Siler, W. A competing-risk model for animal mortality. Ecology 60, 750–757 (1979).

59. Millar, R. B. Comparison of hierarchical bayesian models for overdispersed count data using DIC and Bayes’ factors. Biometrics 65, 962–969 (2009).

60. McCulloch, R. E. Local model influence. J. Am. Stat. Assoc. 84, 473–478 (1989).

61. Kullback, S. & Leibler, R. A. On information and sufficiency. Ann. Math. Stat. 22, 79–86 (1951).

