## Supplementary Material for "Silver-spoon upbringing improves early-life fitness but promotes reproductive ageing in a wild bird"

**Running Title**: Good natal conditions fasten ageing in nature

**Keywords**: life history evolution, disposable soma, reproduction, survival, early-life conditions, ageing, senescence, condition-dependence, trade-offs, brood size manipulation

### Supplementary Tables and Figures

#### Female tarsus length

The tarsus length of female collared flycatchers raised in nests with low-competition (reduced brood) had no difference with the tarsus length of their counterparts raised in nests with high-competition (Fig. S1; $\chi^2 = 0.52$, $P = 0.77$).

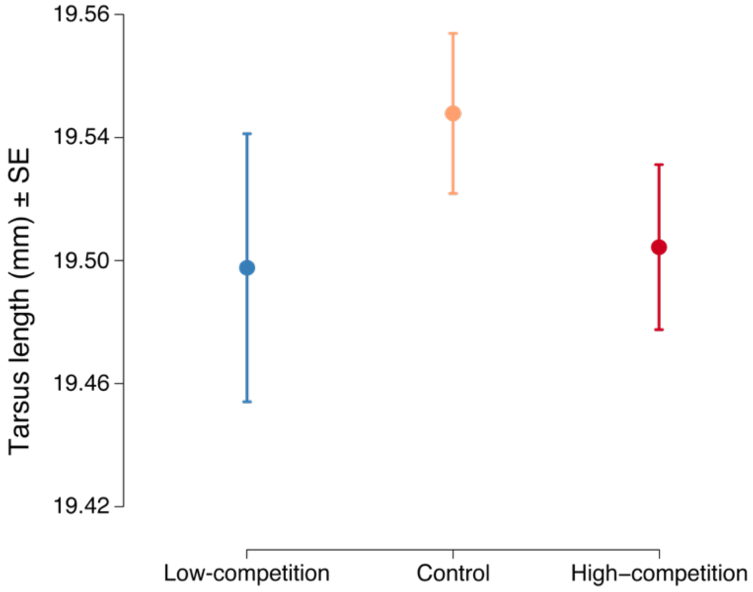


**Figure S1.** Tarsus length (means ± SE) of the three manipulated natal environments. Blue lines represent the increased low-competition treatment, orange lines represent the control treatment and red lines represent the decreased high-competition treatment.

#### Lifetime Reproductive Success

The quality of the natal environment had no effect on lifetime reproduction in the restricted dataset (Fig. 3C, Table S1).

**Table S1.** Generalized linear mixed effects model examining the effect of early life environmental conditions on lifetime reproduction (i.e. total number of recruits over all reproductive events). The reference level for treatment is "Low-competition". Significant effects are in bold.

| **Variable** | **Estimate ± S.E.** | **Z** | ***P*-value (Z)** | **$\chi^2$** | **df** | ***P*-value ($\chi^2$)** | **Variance** |
|---|---|---|---|---|---|---|---|
| **Intercept** | **-0.98 ± 0.26** | **-3.82** | **< 0.001** | **14.60** | **1** | **< 0.001** | |
| Nest Treatment | | | | 1.00 | 2 | 0.607 | |
| Control Trt | -0.15 ± 0.27 | -0.58 | 0.563 | | | | |
| High-competition Trt | -0.28 ± 0.28 | -1.00 | 0.317 | | | | |
| Random | | | | | | | |
| Nest ID | | | | | | | 0.68 |
| Year of Birth | | | | | | | 0.10 |

#### Rate-sensitive Individual Fitness $\lambda_{ind}$

The quality of the natal environment had no effect on individual fitness in the restricted dataset (Fig. 3D, Table S2).

**Table S2.** Linear mixed effects model examining the effect of early life environmental conditions on rate sensitive individual fitness $\lambda_{ind}$. The reference level for treatment is "Low-competition". Significant effects are in bold.

| **Variable** | **Estimate ± S.E.** | **t** | ***P*-value (t)** | **$\chi^2$** | **df** | ***P*-value ($\chi^2$)** | **Variance** |
|---|---|---|---|---|---|---|---|
| **Intercept** | **0.39 ± 0.01** | **40.26** | **< 0.001** | | | | |
| Nest Treatment | | | | 2.17 | 2 | 0.337 | |
| Control Trt | -0.02 ± 0.01 | -1.47 | 0.142 | | | | |
| High-competition Trt | -0.02 ± 0.01 | -1.49 | 0.136 | | | | |
| Random | | | | | | | |
| Nest ID | | | | | | | 0.001 |
| Year of Birth | | | | | | | 0.000 |
| Residual | | | | | | | 0.004 |

#### Age-specific Reproduction

We run 35 different GLMM models with Poisson error structure, to analyse the effects of brood size manipulation on the age-specific reproduction. We first started with a full model, which was backwards reduced with all possible order of term deletion. The models were then compared with the conservative Akaike Information Criterion for small sample size (AIC$_C$) (Table S3). Models 17, 24 and 31 had the best fit and similar outcomes. Therefore, we presented the output of the most comprehensive model (i.e. model 17) in the Results section and the output of the other two models below (Table S4).

We then ran these three best models (i.e. models 17, 24 and 31) using Bayesian Markov-chain Monte Carlo methods, where posterior distributions allow calculation of proper estimate errors. We run the models for 1 200 000 iterations, discarded the first 200 000 iterations as burn-in and used interval sampling every 1000 steps. We used uninformative priors (inverse Wishart with V = 1 and nu = 0.002) for all the models, and assessed model convergence from the low autocorrelation levels (< 0.05) of all parameter estimates and graphically from the posterior estimate diagnostic plots. With this approach we obtained the same results as above (Table S5), and thus we used the *predict* fuction to calculate and plot the model estimates from the frequentist GLMM Poisson model 17 (Figure 3B).

**Table S3.** All generalized linear mixed effects models examining the effect of early life environmental conditions on age-specific reproduction. Model selection was based on the Akaike's Information Criterion adjusted for small sample sizes (AICc) value, delta AICc (ΔAICc) & Akaike weight (wi). The best three models are denoted with bold font.

| Model ID | Model Variables | AIC$_C$ | ΔAIC$_C$ | AIC$_C$Wt |
|---|---|---|---|---|
| **24** | **Trt*Age + Age$^2$ + ALR + AFR + Trt:ALR + Age:AFR + Age$^2$:AFR** | **958.6** | **0.0** | **0.194** |
| **31** | **Trt*Age + Age$^2$ + ALR + AFR + Age:AFR + Age$^2$:AFR** | **958.9** | **0.3** | **0.167** |
| **17** | **Trt*Age + Age$^2$ + ALR + AFR + Trt:ALR + Age:ALR + Age:AFR + Age$^2$:AFR** | **959.5** | **0.9** | **0.124** |
| 32 | Trt*Age + Age$^2$ + ALR + AFR + Trt:ALR + Age:AFR | 961.3 | 2.7 | 0.050 |
| 15 | Trt*Age + Age$^2$ + ALR + AFR + Trt:ALR + Age:ALR + Age$^2$:ALR + Age:AFR + Age$^2$:AFR | 961.4 | 2.8 | 0.048 |

| | | | | |
|---|---|---|---|---|
| 28 | Trt*Age + $Age^2$ + ALR + AFR + Trt:ALR + Age:ALR + Age:AFR | 961.5 | 2.9 | 0.046 |
| 35 | Trt*Age + $Age^2$ + ALR + AFR + Age:AFR | 961.6 | 3.0 | 0.043 |
| 18 | Trt*Age + $Age^2$ + ALR + AFR + Trt:$Age^2$ + Trt:ALR + Age:AFR + $Age^2$:AFR | 962.0 | 3.4 | 0.036 |
| 16 | Trt*Age + $Age^2$ + ALR + AFR + Trt:ALR + Trt:AFR + Age:AFR + $Age^2$:AFR | 962.1 | 3.5 | 0.034 |
| 26 | Trt*Age + $Age^2$ + ALR + AFR + Trt:$Age^2$ + Age:AFR + $Age^2$:AFR | 962.1 | 3.5 | 0.032 |
| 25 | Trt*Age + $Age^2$ + ALR + AFR + Trt:AFR + Age:AFR + $Age^2$:AFR | 962.2 | 3.6 | 0.032 |
| 9 | Trt*Age + $Age^2$ + ALR + AFR + Trt:ALR + Trt:AFR + Age:ALR + Age:AFR + $Age^2$:AFR | 962.9 | 4.3 | 0.023 |
| 19 | Trt*Age + $Age^2$ + ALR + AFR + Trt:$Age^2$ + Trt: AFR + Age:AFR + $Age^2$:AFR | 962.9 | 4.3 | 0.023 |
| 21 | Trt*Age + $Age^2$ + ALR + AFR + Trt:ALR + Age:ALR + $Age^2$:ALR + Age:AFR | 962.9 | 4.3 | 0.023 |
| 11 | Trt*Age + $Age^2$ + ALR + AFR + Trt:$Age^2$ + Trt:ALR + Age:ALR + Age:AFR + $Age^2$:AFR | 963.1 | 4.5 | 0.021 |
| 33 | Trt*Age + $Age^2$ + ALR + AFR + Trt:AFR + Age:AFR | 964.1 | 5.5 | 0.012 |
| 27 | Trt*Age + $Age^2$ + ALR + AFR + Trt:ALR + Trt:AFR + Age:AFR | 964.3 | 5.7 | 0.011 |
| 20 | Trt*Age + $Age^2$ + ALR + AFR + Trt:ALR + Trt:AFR + Age:ALR + Age:AFR | 964.3 | 5.7 | 0.011 |
| 29 | Trt*Age + $Age^2$ + ALR + AFR + Trt:$Age^2$ + Trt:ALR + Age:AFR | 964.6 | 6.0 | 0.010 |
| 8 | Trt*Age + $Age^2$ + ALR + AFR + Trt:$Age^2$ + Trt:ALR + Age:ALR + $Age^2$:ALR + Age:AFR + $Age^2$:AFR | 964.7 | 6.1 | 0.009 |
| 7 | Trt*Age + $Age^2$ + ALR + AFR + Trt:ALR + Trt:AFR + Age:ALR + $Age^2$:ALR + Age:AFR + $Age^2$:AFR | 964.9 | 6.3 | 0.008 |
| 34 | Trt*Age + $Age^2$ + ALR + AFR + Trt:$Age^2$ + Age:AFR | 964.9 | 6.3 | 0.008 |
| 23 | Trt*Age + $Age^2$ + ALR + AFR + Trt:$Age^2$ + Trt:ALR + Age:ALR + Age:AFR | 965.0 | 6.4 | 0.008 |
| 10 | Trt*Age + $Age^2$ + ALR + AFR + Trt:$Age^2$ + Trt:ALR + Trt:AFR + Age:AFR + $Age^2$:AFR | 965.6 | 7.0 | 0.006 |
| 12 | Trt*Age + $Age^2$ + ALR + AFR + Trt:ALR + Trt:AFR + Age:ALR + $Age^2$:ALR + Age:AFR | 966.0 | 7.4 | 0.005 |
| 14 | Trt*Age + $Age^2$ + ALR + AFR + Trt:$Age^2$ + Trt:ALR + Age:ALR + $Age^2$:ALR + Age:AFR | 966.1 | 7.5 | 0.005 |
| 5 | Trt*Age + $Age^2$ + ALR + AFR + Trt:$Age^2$ + Trt:ALR + Trt:AFR + Age:ALR + Age:AFR + $Age^2$:AFR | 966.6 | 8.0 | 0.004 |
| 30 | Trt*Age + $Age^2$ + ALR + AFR + Trt:$Age^2$ + Trt:AFR + Age:AFR | 967.6 | 9.0 | 0.002 |

| | | | | |
|---|---|---|---|---|
| 22 | Trt*Age + Age$^2$ + ALR + AFR + Trt:Age$^2$ + Trt:ALR + Trt: AFR + Age:AFR | 967.7 | 9.1 | 0.002 |
| 13 | Trt*Age + Age$^2$ + ALR + AFR + Trt:Age$^2$ + Trt:ALR + Trt:AFR + Age:ALR + Age:AFR | 967.9 | 9.3 | 0.002 |
| 4 | Trt*Age + Age$^2$ + ALR + AFR + Trt:Age$^2$ + Trt:ALR + Trt:AFR + Age:ALR + Age$^2$:ALR + Age:AFR + Age$^2$:AFR | 968.3 | 9.7 | 0.001 |
| 6 | Trt*Age + Age$^2$ + ALR + AFR + Trt:Age$^2$ + Trt:ALR + Trt:AFR + Age:ALR + Age$^2$:ALR + Age:AFR | 969.3 | 10.7 | 0.001 |
| 3 | Trt*Age*ALR + Age$^2$ + AFR + Trt:Age$^2$ + Trt:AFR + Age$^2$:ALR + Age:AFR + Age$^2$:AFR | 970.6 | 12.0 | 0.001 |
| 2 | Trt*Age*ALR + Age$^2$ + AFR + Trt:Age$^2$ + Trt:AFR + Age$^2$:ALR + Age:AFR + Age$^2$:AFR + Trt:Age:AFR | 973.3 | 14.7 | 0.000 |
| 1 | Trt*Age*ALR + Age$^2$ + AFR + Trt:Age$^2$ + Trt:AFR + Age$^2$:ALR + Age:AFR + Age$^2$:AFR + Trt:Age$^2$:ALR | 974.5 | 15.9 | 0.000 |
| Full | Trt*Age*ALR + Age$^2$ + AFR + Trt:Age$^2$ + Trt:AFR + Age$^2$:ALR + Age:AFR + Age$^2$:AFR + Trt:Age:AFR + Trt:Age$^2$:ALR | 976.6 | 18.0 | 0.000 |

**Table S4.** The other two best generalized linear mixed effects models examining the effect of early life environmental conditions on age-specific reproduction (Model 24 and Model 31). Significant effects are in bold.

| Variable | Estimate ± S.E. | Z | *P*-value (Z) | $\chi^2$ | df | *P*-value ($\chi^2$) | Variance |
|---|---|---|---|---|---|---|---|
| *Model 24* | | | | | | | |
| **Intercept** | **-1.65 ± 0.26** | **-6.37** | **< 0.001** | **40.56** | **1** | **< 0.001** | |
| **Age** | **0.66 ± 0.27** | **2.44** | **0.015** | **5.97** | **1** | **0.015** | |
| **Age$^2$** | **-0.43 ± 0.12** | **-3.72** | **< 0.001** | **13.82** | **1** | **< 0.001** | |
| **Age at First Repr. (AFR)** | **-0.86 ± 0.16** | **-5.34** | **< 0.001** | **28.55** | **1** | **< 0.001** | |
| Age at Last Repr. (ALR) | 0.27 ± 0.19 | 1.46 | 0.145 | 2.13 | 1 | 0.145 | |
| Nest Treatment | | | | 2.67 | 2 | 0.263 | |
| Control Trt | -0.30 ± 0.21 | -1.39 | 0.163 | | | | |
| High-competition Trt | -0.31 ± 0.21 | -1.45 | 0.147 | | | | |
| **Interaction (Age × AFR)** | **0.95 ± 0.24** | **3.97** | **< 0.001** | **15.74** | **1** | **< 0.001** | |
| **Interaction (Age$^2$ × AFR)** | **-0.23 ± 0.11** | **-1.99** | **0.047** | **3.95** | **1** | **0.047** | |

| | | | | | | | |
|---|---|---|---|---|---|---|---|
| **Interaction (Age × Trt)** | | | | **7.85** | **2** | **0.020** | |
| Age x Control Trt | 0.21 ± 0.29 | 0.72 | 0.471 | | | | |
| **Age × High-competition Trt** | **0.75 ± 0.28** | **2.65** | **0.008** | | | | |
| Interaction (ALR × Trt) | | | | 4.58 | 2 | 0.101 | |
| **ALR × Control Trt** | **-0.53 ± 0.26** | **-2.02** | **0.043** | | | | |
| ALR × High-competition Trt | -0.41 ± 0.25 | -1.65 | 0.099 | | | | |
| Random | | | | | | | |
| Female ID | | | | | | | 0.00 |
| Nest ID | | | | | | | 0.24 |
| Year of Birth | | | | | | | 0.00 |
| Year of Annual Repr. | | | | | | | 0.42 |
| Observation | | | | | | | 0.22 |
| *Model 31* | | | | | | | |
| **Intercept** | **-1.68 ± 0.26** | **-6.45** | **< 0.001** | **41.57** | **1** | **< 0.001** | |
| **Age** | **0.83 ± 0.26** | **3.18** | **0.001** | **10.09** | **1** | **0.001** | |
| **$Age^2$** | **-0.43 ± 0.12** | **-3.68** | **< 0.001** | **13.51** | **1** | **< 0.001** | |
| **Age at First Repr. (AFR)** | **-0.87 ± 0.16** | **-5.41** | **< 0.001** | **29.25** | **1** | **< 0.001** | |
| Age at Last Repr. (ALR) | -0.06 ± 0.10 | -0.63 | 0.528 | 0.40 | 1 | 0.528 | |
| Nest Treatment | | | | 2.19 | 2 | 0.335 | |
| Control Trt | -0.26 ± 0.21 | -1.21 | 0.228 | | | | |
| High-competition Trt | -0.29 ± 0.21 | -1.36 | 0.173 | | | | |
| **Interaction (Age × AFR)** | **0.95 ± 0.24** | **3.99** | **< 0.001** | **15.95** | **1** | **< 0.001** | |
| **Interaction ($Age^2$ × AFR)** | **-0.23 ± 0.11** | **-1.98** | **0.047** | **3.94** | **1** | **0.047** | |
| **Interaction (Age × Trt)** | | | | **9.71** | **2** | **0.008** | |
| Age × Control Trt | -0.09 ± 0.25 | -0.37 | 0.709 | | | | |
| **Age × High-competition Trt** | **0.53 ± 0.24** | **2.18** | **0.029** | | | | |
| Random | | | | | | | |
| Female ID | | | | | | | 0.00 |
| Nest ID | | | | | | | 0.26 |
| Year of Birth | | | | | | | 0.00 |
| Year of Annual Repr. | | | | | | | 0.41 |
| Observation | | | | | | | 0.24 |

**Table S5.** The three best models run with a Bayesian framework examining the effect of early life environmental conditions on age-specific reproduction. The results are the same with the frequentist models 17, 24 and 31 (Table 1 and Table S4). Significant effects are in bold.

| **Variable** | **Posterior Mean** | **Lower 95% C.I.** | **Higher 95% C.I.** | ***P*-value** |
|---|---|---|---|---|
| *Model 17* | | | | |
| **Intercept** | **-1.72** | **-2.29** | **-1.13** | **< 0.001** |
| **Age** | **0.67** | **0.14** | **1.16** | **0.012** |
| **Age$^2$** | **-0.59** | **-0.91** | **-0.32** | **< 0.001** |
| **Age at First Repr. (AFR)** | **-0.89** | **-1.19** | **-0.57** | **< 0.001** |
| Age at Last Repr. (ALR) | 0.33 | -0.04 | 0.70 | 0.088 |
| Nest Treatment | | | | |
| Control Trt | -0.31 | -0.72 | 0.11 | 0.158 |
| High-competition Trt | -0.33 | -0.74 | 0.08 | 0.122 |
| Interaction (Age × ALR) | 0.17 | -0.12 | 0.42 | 0.220 |
| **Interaction (Age × AFR)** | **0.97** | **0.49** | **1.44** | **< 0.001** |
| **Interaction (Age$^2$ × AFR)** | **-0.22** | **-0.41** | **0.00** | **0.036** |
| Interaction (Age × Trt) | | | | |
| Age × Control Trt | 0.24 | -0.33 | 0.81 | 0.426 |
| **Age × High-competition Trt** | **0.81** | **0.21** | **1.32** | **0.002** |
| Interaction (ALR × Trt) | | | | |
| **ALR × Control Trt** | **-0.59** | **-1.08** | **-0.05** | **0.032** |
| **ALR × High-competition Trt** | **-0.51** | **-1.04** | **-0.02** | **0.044** |
| Random | | | | |
| Female ID | 0.08 | 0.00 | 0.39 | |
| Nest ID | 0.23 | 0.00 | 0.56 | |
| Year of Birth | 0.07 | 0.00 | 0.29 | |
| Year of Annual Repr. | 0.60 | 0.00 | 1.30 | |
| Residual variance | 0.18 | 0.00 | 0.64 | |
| *Model 24* | | | | |
| **Intercept** | **-1.76** | **-2.33** | **-1.19** | **< 0.001** |
| **Age** | **0.72** | **0.23** | **1.26** | **0.006** |
| **Age$^2$** | **-0.49** | **-0.70** | **-0.25** | **< 0.001** |
| **Age at First Repr. (AFR)** | **-0.91** | **-1.25** | **-0.64** | **< 0.001** |
| Age at Last Repr. (ALR) | 0.28 | -0.07 | 0.633 | 0.118 |

| | | | | |
|---|---|---|---|---|
| Nest Treatment | | | | |
| Control Trt | -0.28 | -0.71 | 0.21 | 0.190 |
| High-competition Trt | -0.28 | -0.68 | 0.18 | 0.202 |
| **Interaction (Age x AFR)** | **1.01** | **0.54** | **1.47** | **< 0.001** |
| **Interaction (Age$^2$ x AFR)** | **-0.24** | **-0.48** | **-0.05** | **0.016** |
| Interaction (Age x Trt) | | | | |
| Age x Control Trt | 0.19 | -0.39 | 0.76 | 0.560 |
| **Age x High-competition Trt** | **0.75** | **0.21** | **1.27** | **0.002** |
| Interaction (ALR x Trt) | | | | |
| **ALR x Control Trt** | **-0.56** | **-1.11** | **-0.05** | **0.034** |
| ALR x High-competition Trt | -0.42 | -0.89 | 0.05 | 0.078 |
| Random | | | | |
| Female ID | 0.07 | 0.00 | 0.28 | |
| Nest ID | 0.23 | 0.00 | 0.61 | |
| Year of Birth | 0.06 | 0.00 | 0.62 | |
| Year of Annual Repr. | 0.66 | 0.05 | 1.44 | |
| Residual variance | 0.18 | 0.00 | 0.62 | |
| *Model 31* | | | | |
| **Intercept** | **-1.74** | **-2.31** | **-1.21** | **< 0.001** |
| **Age** | **0.82** | **0.31** | **1.36** | **0.002** |
| **Age$^2$** | **-0.48** | **-0.75** | **-0.24** | **< 0.001** |
| **Age at First Repr. (AFR)** | **-0.90** | **-1.19** | **-0.55** | **< 0.001** |
| Age at Last Repr. (ALR) | -0.06 | -0.27 | 0.15 | 0.568 |
| Nest Treatment | | | | |
| Control Trt | -0.23 | -0.67 | 0.17 | 0.308 |
| High-competition Trt | -0.29 | -0.68 | 0.11 | 0.158 |
| **Interaction (Age x AFR)** | **1.01** | **0.54** | **1.57** | **< 0.001** |
| **Interaction (Age$^2$ x AFR)** | **-0.25** | **-0.50** | **-0.02** | **0.016** |
| **Interaction (Age x Trt)** | | | | |
| Age x Control Trt | -0.07 | -0.55 | 0.39 | 0.748 |
| **Age x High-competition Trt** | **0.58** | **0.09** | **1.01** | **0.018** |
| Random | | | | |
| Female ID | 0.09 | 0.00 | 0.42 | |
| Nest ID | 0.22 | 0.00 | 0.59 | |
| Year of Birth | 0.07 | 0.00 | 0.30 | |
| Year of Annual Repr. | 0.60 | 0.00 | 1.32 | |
| Residual variance | 0.21 | 0.00 | 0.65 | |

#### Survival & Mortality rates

We first explored the age-specific mortality rates with models following the Weibull, Gompertz, Logistic and Exponential distributions and then compared them with the Deviance Information Criterion (DIC) (Table S7). The model with the most appropriate mortality function for our data was the Logistic model with bathtub shape (Table S6).

**Table S6.** Parameter estimates for each treatment from the Logistic model with bathtub shape (best fitting model). LC: Low Competition, HC: High Competition

| Parameter | Treatment | Estimates | Lower 95% CI | Upper 95% CI | Ser. autocor |
|---|---|---|---|---|---|
| $\alpha_0$ | Control | -4.144 | -5.453 | -3.073 | -0.040 |
| | Decreased – LC | -3.775 | -5.138 | -2.656 | 0.019 |
| | Increased – HC | -4.053 | -5.364 | -2.890 | -0.005 |
| $\alpha_1$ | Control | 1.158 | 0.084 | 2.725 | -0.013 |
| | Decreased – LC | 1.072 | 0.070 | 2.657 | -0.028 |
| | Increased – HC | 1.184 | 0.084 | 2.848 | 0.037 |
| c | Control | 0.006 | 0.000 | 0.023 | 0.005 |
| | Decreased – LC | 0.013 | 0.000 | 0.045 | 0.026 |
| | Increased – HC | 0.007 | 0.000 | 0.026 | 0.039 |
| $b_0$ | Control | -4.542 | -5.497 | -3.707 | 0.008 |
| | Decreased – LC | -4.142 | -5.157 | -3.197 | -0.010 |
| | Increased – HC | -4.524 | -5.514 | -3.636 | -0.033 |
| $b_1$ | Control | 2.078 | 1.544 | 2.640 | 0.017 |
| | Decreased – LC | 1.580 | 1.066 | 2.166 | -0.015 |
| | Increased – HC | 2.145 | 1.593 | 2.784 | -0.020 |
| $b_2$ | Control | 2.478 | 1.586 | 3.424 | 0.002 |
| | Decreased – LC | 1.370 | 0.585 | 2.462 | -0.005 |
| | Increased – HC | 2.328 | 1.515 | 3.298 | -0.024 |
| Pr | - | 0.687 | 0.657 | 0.718 | 0.001 |

**Table S7.** Model comparison based on the Deviance Information Criterion (DIC) between ten models we tested with the BaSTA analysis. K indicates the effective number of parameters.

| Model | Shape | K | DIC | ΔDIC |
|---|---|---|---|---|
| Logistic | Bathtub | 19 | 4017.27 | 0 |
| Logistic | Simple | 10 | 4067.97 | 50.70 |
| Logistic | Makeham | 13 | 4081.69 | 64.42 |
| Weibull | Bathtub | 16 | 4141.46 | 124.19 |
| Weibull | Simple | 7 | 4188.79 | 171.52 |
| Weibull | Makeham | 10 | 4206.49 | 189.22 |
| Gompertz | Bathtub | 16 | 4452.32 | 435.05 |
| Gompertz | Simple | 7 | 4516.62 | 499.35 |
| Gompertz | Makeham | 10 | 4530.57 | 513.30 |
| Exponential | Simple | 4 | 5088.74 | 1071.47 |
